# Political legacies shape bear use of anthropogenic spaces

**DOI:** 10.1101/2025.11.20.689425

**Authors:** Catalina Munteanu, Mihai Daniel Nita, Ana Stritih, Simone Roverelli, Benjamin Michael Kraemer, Ovidiu Ionescu, Ilse Storch

## Abstract

Romania hosts Europe’s largest brown bear population west of Russia, but rising human-bear encounters and related casualties raise concerns about their drivers and future population management. Using >10,000 encounter reports (2019–2024) in a hierarchical occupancy model accounting for detection processes, we show that bear presence in human-dominated spaces is linked to food conditioning, habituation, and shaped by historical management. Bears occur more often in high wildland-urban interface areas, especially those closer to communist-era elite-hunting grounds with feeding sites. These patterns reflect a long-term legacy of 1960s population management policies interacting with environmental and behavioral processes to shape recent bear behavior. We call for management addressing ecological and social sustainability by phasing out supplemental feeding, reducing habituation through a restored landscape of fear, improving public knowledge of bear ecology, and recognizing historical legacies to promote coexistence.

**Highlights:**

- We used hierarchical species occupancy models to analyze brown bears’ use of human-dominated spaces in Romania.
- Bear detectability depends strongly on activity patterns of both bears and humans
- Bear occurrence in anthropogenic spaces is strongly associated with high wildland–urban interface areas and abundant anthropogenic food waste.
- Bear occurrence in WUI areas is modulated by historical population management legacies.

## Introduction

As global human population grows (Taagepera and Nemčok 2024; Cohen 1995), pressure on natural environments intensifies, especially in densely populated regions like Europe (Weber and Sciubba 2019). Despite increasing anthropogenic pressure, Europe retains large wilderness areas like the Romanian Carpathians, home to the largest brown bear (Ursus arctos) population west of Russia - estimated at 10000-12000 individuals - alongside other iconic large mammals (Chapron et al. 2014; Mráz and Ronikier 2016; Dyck et al. 2022). With a habitat comparable in size to the Greater Yellowstone Ecosystem, the region holds over 15 times more brown bears (Quammen 2003). This abundance results from long-term habitat suitability, historic and recent management, and greater social tolerance toward large carnivores compared to Central Europe (Cristescu et al. 2019; Rode et al. 2021). However, in recent decades, reports of human-bear interactions—including livestock depredation and attacks in urban or peri-urban areas—have increased (Pop, Gradinaru, et al. 2023; Pop, Dyck, et al. 2023; Bombieri et al. 2019), raising questions about human-bear coexistence. Amid increasing anthropogenic resistance (Ghoddousi et al. 2021) and declining tolerance, two competing management strategies emerge: reconnecting habitats to support growing populations and promote coexistence (Mayer et al. 2023; Fletcher and Toncheva 2021; Can et al. 2014) and or using hunting to reduce numbers, restore fear, limit habituation, and fund conservation (Wheat and Wilmers 2016; L. Brown et al. 2023; Hertel et al. 2016; Stăncioiu, Dutcă, et al. 2019). The success of either approach depends on understanding the socio-ecological drivers of increasing urban and peri-urban bear encounters, and designing and adopting bear management strategies policies that balance conservation success with human safety and inter-specific coexistence.

The recent recovery of large mammal populations is not unique to Romania; many developed countries have seen similar trends due to legal protection and coexistence efforts (Chapron et al. 2014). However, the intense use of urban and peri-urban habitats by bears is unique to Romania, providing an unprecedented natural experiment on how long-term protection and management can interact to shape ecological responses (Fig 1). As adaptable omnivores with ranges up to 800 km^2^, brown bears typically inhabit extended forested habitats, but can successfully exploit anthropogenic food sources and urban peripheries (Lucas et al. 2025). Their growing urban presence reflects ecological and behavioral responses to human exposure and environmental pressures: (Elfström et al. 2014). The shift towards increased use of anthropogenic spaces can be explained by several mechanisms, such as habitat fragmentation, intraspecific competition and displacement of young bears or opportunistic behaviour such as human habituation, food-conditioning (Elfström et al. 2014). Importantly, hunting maintains a landscape of fear, discouraging foraging in proximity of humans (Lodberg-Holm et al. 2019; Gaynor et al. 2019). However, if bears associate humans with food sources and loose fear (Gaynor et al. 2019) they can more often cause property damage, livestock losses, or attack humans (Pop, Dyck et al. 2023; Pop, Grădinaru et al. 2023a), fueling intolerance and fear (Stăncioiu, Dutcă, et al. 2019). Despite recognized trade-offs between food availability and risk-avoidance, their interactions and the time-scales at which the species respond remain widely unexplored.

**Figure 1.**
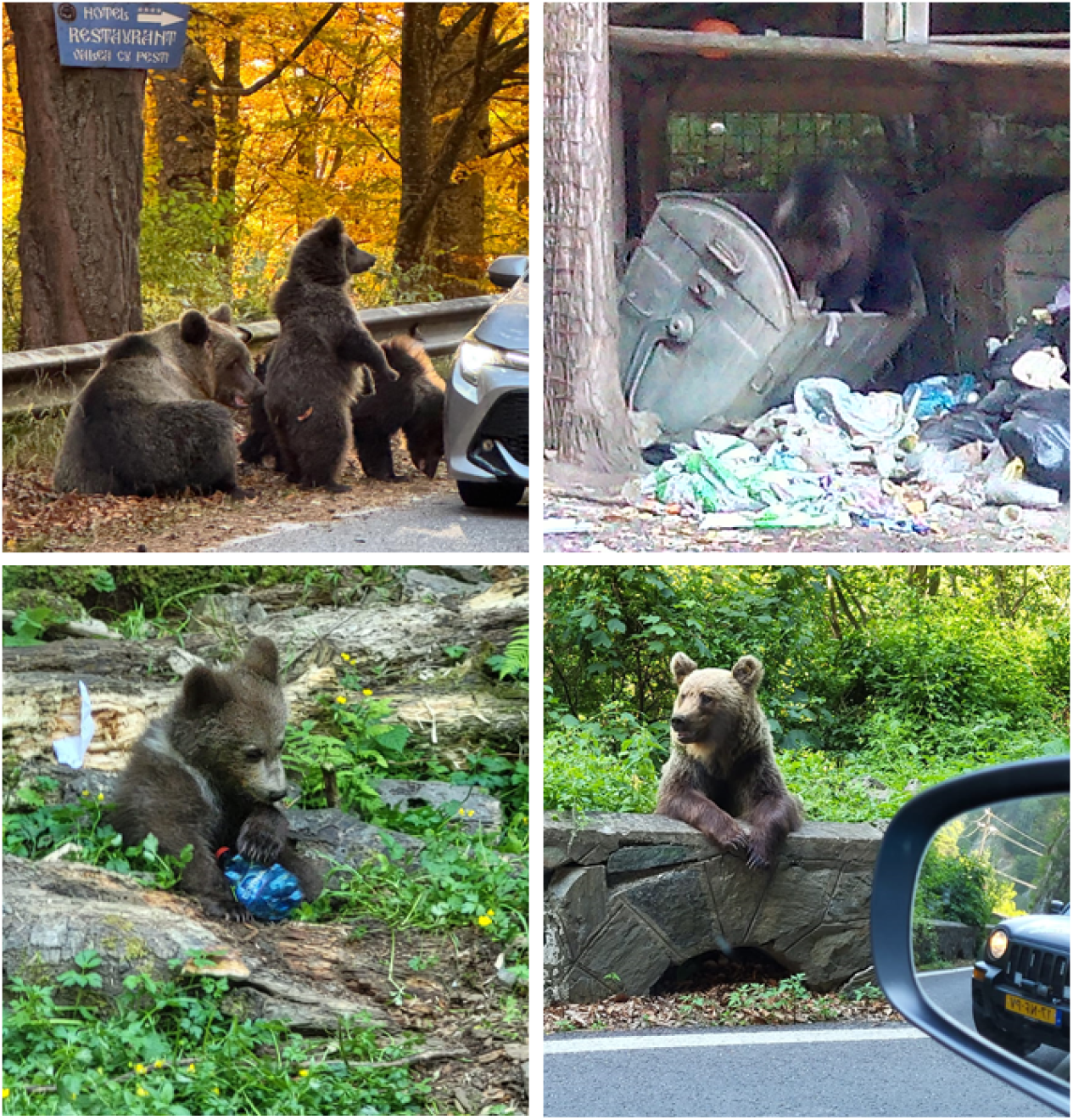
Examples of bears exploiting human-dominated landscapes, supporting the conditioning and habituation hypothesis A. in proximity of restaurants (Foto: M. Nita) B. at garbage dumps (Foto: S Munteanu) C. in Wildland-Urban-Interface (Foto: C Munteanu) and D along roads (Foto: C Munteanu)

Social-ecological systems and species respond to land use and management decisions with time-delays, and impacts can persist from years to centuries (Polaina et al. 2019; Munteanu et al. 2020; Foster et al. 2003, Meyers et al 2025). Ecological legacies can be particularly strong after shock-events like wars (Barthelme et al. 2024; Bragina et al. 2018) or under totalitarian regimes, where centralized environmental interventions served political agendas (Brain 2010). For instance, heavy post–World War II forest harvesting to pay reparations altered forest structures, canopy loss and seed bank diversity in modern Romania and Germany (Nita et al. 2018; Munteanu et al. 2015, Meyers et al 2025). Ecological legacies can span decades to centuries ( Foster et al. 2003, Munteanu et al. 2015, Munteanu et al. 2020)and interact with social legacies, including institutions, norms, and customs that shape management (Tappeiner et al. 2021). Ignoring legacy effects risks misinformed or misguided management and conservation decisions (Munteanu et al. 2020; Stăncioiu et al. 2021).

In Romania, legacies related to bear populations may stem from communist central planning (1947–1989), which imposed strict wildlife monitoring (since 1958) and management plans emphasizing trophy species like bear, wild boar and red deer due to leader Nicolae Ceauşescu’s hunting interests (Salvatori et al. 2002). Bears were protected, but intensively managed, including supplemental feeding at hunting locations since 1974, protected buffer zones around hunting grounds, breeding programs and intensive monitoring (Mooney 1991; Georgescu 2003; Crisan 2010). Bear protection policies were initiated in 1953 when den hunting was banned, and together with communist population management, likely contributed to Romania being a European bear-abundance hotspot (Quammen 2003). Following the communist regime collapse in 1990, supplemental feeding continued in historical locations and also expanded across the species range (at least 2km away from forest edge), while hunting resumed under new regulations, stabilizing populations for several years (Fig 2). When hunting was later restricted under EU law, except for damage-related “intervention culls” (Stăncioiu, Micu, et al. 2019), the landscape of fear likely dissolved (Swenson 1999; Ordiz et al. 2012). As fear of humans decreased, bears increasingly used anthropogenic habitats (Cimpoca and Voiculescu 2022; Swenson 1999) — agricultural fields, farms, orchards, and food waste sites— because they provided reliable food sources (Bautista et al. 2017) and refuge from older dominant males (Bellemain et al. 2005; Steyaert et al. 2012). While political discourse and decisions revolve around hunting, supplemental feeding for bears was active in Romania between 1974 and 2016 and continues in Romania for non-target species, raising the question of whether this practice left behind behavioral and ecological legacies that complicate current coexistence efforts.

**Figure 2.**
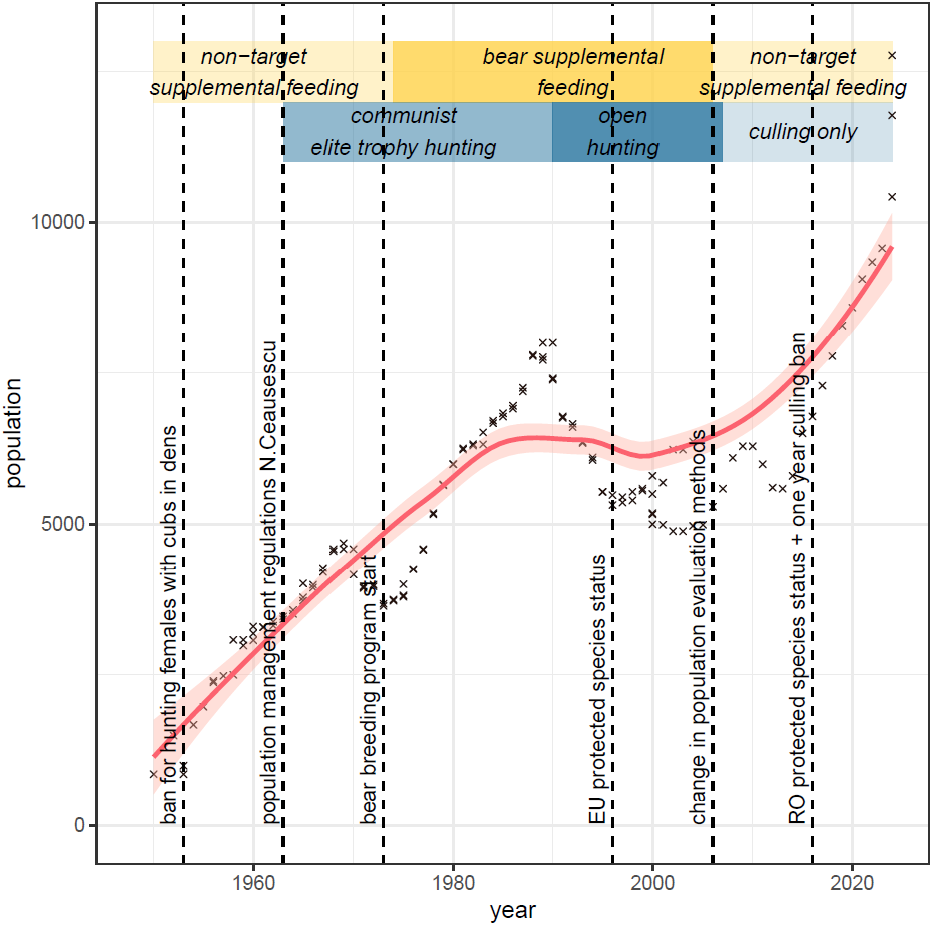
Bear population trends, as compiled from multiple data sources for Romania between 1940 and 2024 in relation to major political decisions regarding population management. Hunting and supplemental feeding regulations are shown in blue and yellow respectively. The trendline is computed using a LOESS smoother with a span of 0.85 and 95% confidence interval. Data sources: Mooney 1991, Quammen 2003, MMP 2024, INCDS 2021, Ionescu 2016.

Our goal was to assess if historical population management affects contemporary human-wildlife interactions,particularly the bears’ use of urban and peri-urban spaces. Specifically, we asked a) where and when bear encounters are most likely in Romania b) what ecological mechanisms drive the use of urban habitats and c) whether contemporary encounter patterns reflect historical management legacies?

To address these questions, we used publicly available, citizen-driven data on bear encounters in Romania (2019-2024). Encounters were reported via emergency line calls and curated by a national text-message warning system (RO-ALERT). Citizen science data on human-wildlife encounters, though spatially and temporally biased, provide valuable insights into species space use. We applied a hierarchical species occupancy model separating observational from ecological processes, to disentangle drivers of bear occurrence in human areas from detection biases. Calls were analyzed in a 3×3 km national grid, modeling detection as a function of time, season, and human presence (tourism, population, road density). Once accounting for detection probability, occurrence was modelled based on population density, habitat structures (forest, wildland-urban-interface: WUI), human food attractants (human food waste), and historical management legacies (distance to historical feeding and hunting grounds). We expected higher detection during crepuscular hours, early spring after emergence from winter sleep, and in populated or accessible areas. Bears were expected to use anthropogenic spaces where buildings are to a high degree intermixed with forest (high WUI, Schug et al. 2023), where there is more food waste (food conditioning hypothesis) (Elfström et al. 2014) and in areas with high bear densities in the core habitat (despotic distribution hypothesis (Elfström et al. 2014). Distance to historical elite-hunting districts—once characterized by high densities, strictly enforced supplemental feeding and bait hunting —was used to test for legacy effects.

We hypothesized that historical management still shapes bear behavior: earlier hunting may have maintained a landscape of fear reducing use of anthropogenic space, whereas continued feeding after hunting bans likely led to habituation and greater human–bear conflict. Romania’s political and environmental history, and its growing population of brown bears offers a unique natural experiment for understanding how management drives long term human–wildlife coexistence - a challenge that many countries with growing large mammal populations face today (Fletcher and Toncheva 2021; Can et al. 2014; Dickman et al. 2011).

## Results

### Where and when bear encounters are most likely

We analyzed N=10472 bear alert messages ( 1-443 calls per 3x3 km grid cell). Bears were most often observed during crepuscular hours, which coincides with feeding times (Fig 3) but amongst crepuscular time encounters were more common in evening and afternoon hours when both bears and humans are active (Fig 3). Reports peaked in late summer-early fall, aligning with hyperphagia when bears accumulate resources prior to hibernation. We identified 1533 grid cells where bears were reported at least once and treated these as positive detections:the detection model showed a positive effect of time of day and day of year on the probability of bear detection (M: 0,33, 95% CI: 0,28-0.38, respectively M: M:0.28, 95%CI: 0.23-0.34). Road density (M:0.16, 95%CI: 0.07-0.24), the density of tourist attractions (restaurants, hotels, M: 0.15, 95%CI: 0.02-0.19), and population density (M: 0.13, 95%CI: 0.04-0.21) significantly increase detectability. In short, bears are most often observed on summer evenings, along roads and in the proximity of tourist attractions.

**Figure 3.**
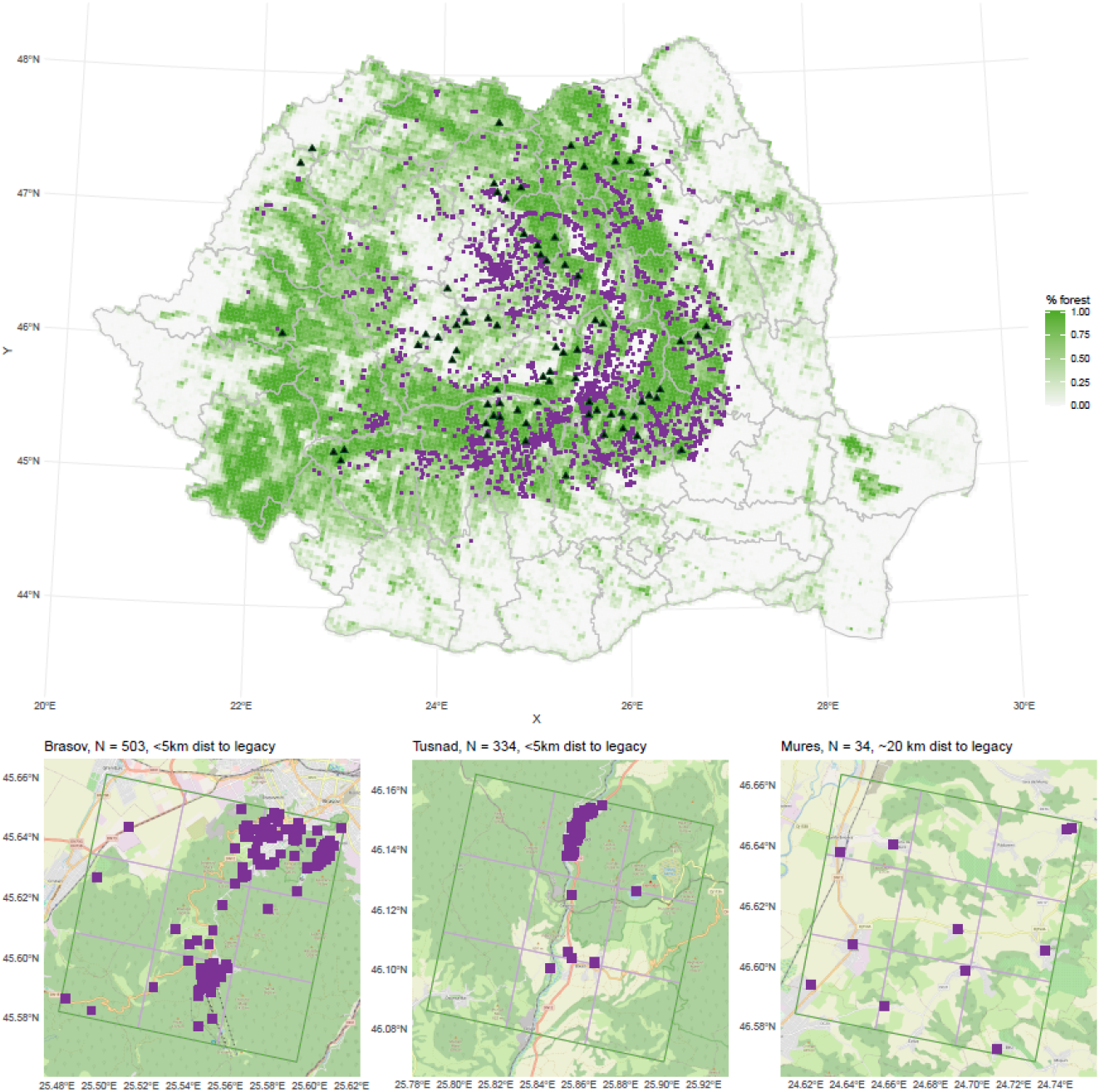
Study area. Romania, with its forest cover and the distribution of human-bear encounters. Each purple point represents a 3x3km grid cell where bear encounters have been reported at least once between 2019-2024 (N=1548 cells, N=10472 calls, range: 1:443 calls/ grid cell). The entire country of Romania is covered by a total of 27197 grid cells. The black triangles represent approximate locations of historical bear feeding and hunting areas.1963-1984. B. Distribution of bear-encounter calls in three subset regions a 9x9 km (point locations may overlap). N represents the number of calls for the entire 9x9 km grid cell. The listed distance is the distance from the cell center to the nearest historical feeding and hunting district.

### Ecological mechanisms driving the use of urban and peri-urban habitats

We found that the percentage of wildland-urban interface (WUI) was the strongest predictor of bear occurrence in urban and peri-urban areas (M:2.87 95%CI: 2.32-3.72). Bears were more likely to occur in WUI cells closer to historically elite hunting grounds - indicating that the proximity to these areas modulates contemporary space-use-patterns of bears (interaction term (M:-0.58, 95%CI: -0.98--0.24). Bear occurrence also increased with the number of anthropogenic food-waste sites per cell (M: 0.61, 95%CI:0.35-0.91), supporting the food-conditioning hypothesis. In contrast, bear density within a 9×9 km area had no significant effect (M = 0.06, 95% CI: –0.28 to 0.43), providing no evidence for the despotic distribution hypothesis. Occurrence probability was higher when forest cover was low or intermediate (M: -0.57, 95%CI: -0.91--0.27). The distance to historical elite-hunting districts had no significant effect on occurrence alone (M: 0.07, 95%CI: -0.50 - 0.66), but only in an interactive effect with WUI. Overall, bears preferentially used human-dominated areas with mixed habitat and abundant food waste, especially near historical hunting and feeding sites.

### Contemporary space use is modulated by historical management legacies

Our hierarchical occupancy models suggest that, after accounting for detectability, bears are more likely to use anthropogenic dominated areas where housing is interspersed with wildland habitat. This relationship is further modulated by proximity to historical elite-hunting districts where supplemental feeding was common: o bear occupancy is higher in WUI areas nearer to these sites (interaction term M:-0.58, 95%CI: -0.98--0.24), suggesting a long-term food conditioning effect. Supporting this, the number of bear-alerts per grid cell was significantly higher near former feeding grounds (R=-0.16 p <0.001, Figure 5). Together, these results point towards a persistent legacy of historical bear management, observed in contemporary bear space use.

## Discussion

Our study shows that both contemporary habitat conditions (high WUI) and historical management legacies (hunting and supplemental feeding regulations) continue to shape human-bear encounters in Romania. Yet, these long-term socio-ecological dynamics remain largely overlooked in recent bear conservation and management debates (Stăncioiu, Dutcă, et al. 2019; Can et al. 2014; Braczkowski et al. 2023). A key sustainability challenge is balancing human wellbeing with biodiversity protection and ecosystem service provision (Braczkowski et al. 2023). The Romanian case provides valuable insights into how politically driven decisions, if disregarded, may have long term impacts on species and their interaction with humans. Decisions made over 50 years ago still influence present human–bear interactions, especially in high WUI areas near former elite-hunting sites. Furthermore, our results provide compelling evidence for the food conditioning and the habituation hypothesis, showing that anthropogenic food sources (proxied by waste locations and habitat intermix) drive bear use of urban and peri-urban areas. These results hold after accounting for detection biases—encounters being most frequent in summer evenings, near roads, and tourist sites, where tourists often report bear encounters. Taken together, our results stress the need to make far-sighted conservation and management decisions today, as today’s actions may shape wildlife behavior for decades.

### Ecological mechanisms and legacy effects

Our results confirm that bears in Romania effectively exploit human-dominated landscapes for food, supporting the habituation and food-conditioning hypotheses. Globally, food-related habituation is a key driver of human-bear interactions (Smith et al. 2005; Elfström et al. 2014; Powell et al. 2022; Carter and Linnell 2016). In Romania, food conditioning appears to be linked to two potential food sources: orchards, agricultural fields (esp. corn) and small farms for fruit and honey (proxied by WUI) and anthropogenic food waste (proxied by garbage dumps, restaurants and hotels). Similar to findings in Poland, bears frequent WUI areas to find fruit, honey and occasionally livestock (Kaim et al. 2025). Indeed, WUI is also used by other large carnivores globally (Odden et al. 2014; Blecha et al. 2018). Over time, anthropogenic food sources lead to food conditioning and habituation behaviours (Kaim et al. 2025). Besides WUI, food waste abundance was a major predictor of bear occurrence. Although waste-management policies exist since 2003, poor infrastructure, coordination, and awareness limit implementation (Nastase et al. 2019). Consequently, bears and related conflicts are most frequent near restaurants, hotels and other food-waste sites (Fig 1,4). Overall, the most important drivers of bear occurrence in human-dominated landscapes are related to food availability - and their long term persistence (Kaim et al. 2025).In Romania, bears have long been exposed to human-related food since targeted supplemental feeding began in the 1974, driven partly by the authoritarian regime’s management practices. Indeed, many authoritarian regimes display political supremacy through control of nature (Brain and Pál 2018; Brain 2010). To display political supremacy through international trophy hunting, Nicolae Ceauşescu implemented a program combining elite hunting, large-scale supplemental feeding (carcasses, pellets, fruit, sweets), and bear breeding and translocation (Nowak et al. 2014; Quammen 2003; Mooney 1991).These measures rapidly boosted populations—from about 1,000 bears in the 1950s to 8,000 by 1989—despite ∼450 individuals hunted by elites (Fig 2) (Georgescu 2003; Crisan 2010). Here, we use the centroids of historically hunting districts as proxies for historical hunting and feeding locations, acknowledging uncertainty since district size (≈100 km^2^) likely obscures true feeding locations. Furthermore, supplemental feeding persisted at these locations and even extended beyond them (but no accurate data on more recent feeding locations was available to us). Our analyses indicate greater bear use of WUI areas near historical hunting districts, suggesting decades-long habituation, but we note estimates of feeding locations are both approximate and likely conservative. The effects may also be partially confounded by the fact that historical elite-hunting grounds were typically established in areas with already high baseline bear populations. Due to these limitations, we expect that our estimates of the effect of the supplemental feeding sites is rather conservative. Incorporating historical population data, historical bear density estimates and updated feeding-site locations (which anecdotally increased threefold between 1990 and 2016) would likely strengthen model explanatory power and increase estimated effect sizes.

**Figure 4.**
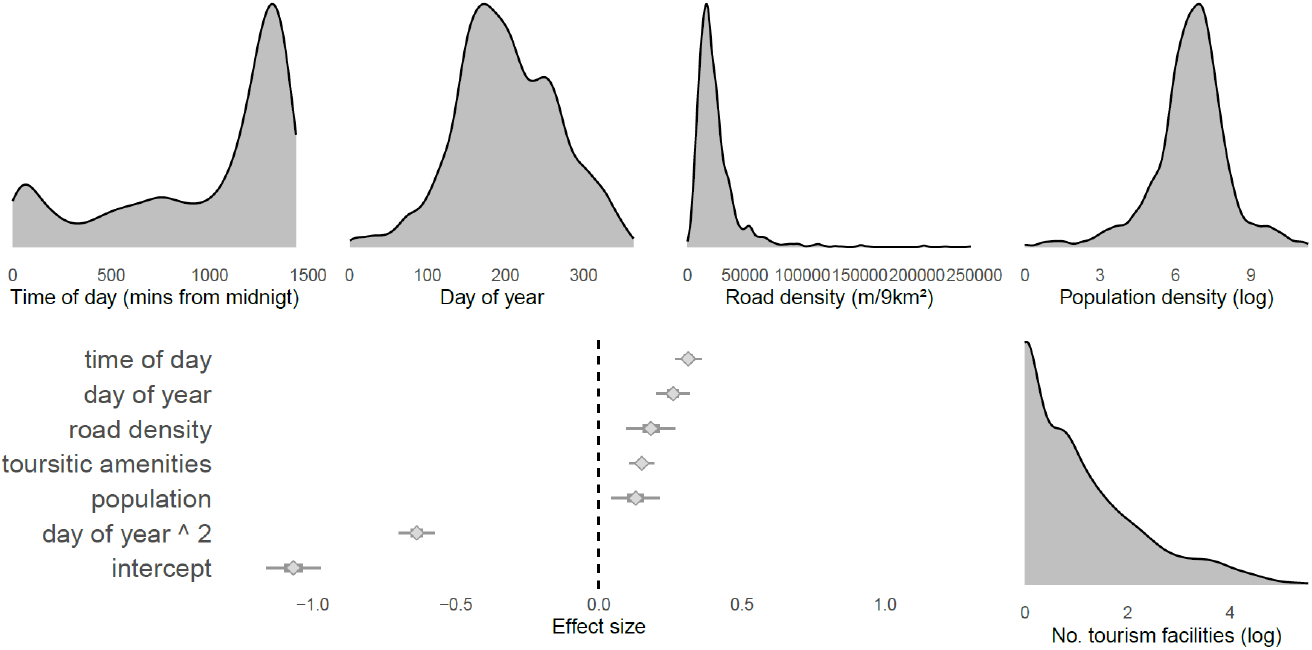
Effect size (posterior mean, 50 and 95 CI) of 5 potential proxies for bear detectability by humans. Detection can be affected by bear activity patterns (tod:time of day, doy:day of year) as well as by human activity and presence (population, number of tourist attractions, density of roads). All variables have a large effect on encounters, which are more likely later in the day, later in the year in areas with higher road density and more touristic attractions. The doy has a humped-shaped effect with encounters more likely to occur in late summer. Grey histograms represent the distributions of all (N=10472) bear calls in Romania recorded between 2019-2024.

**Figure 5.**
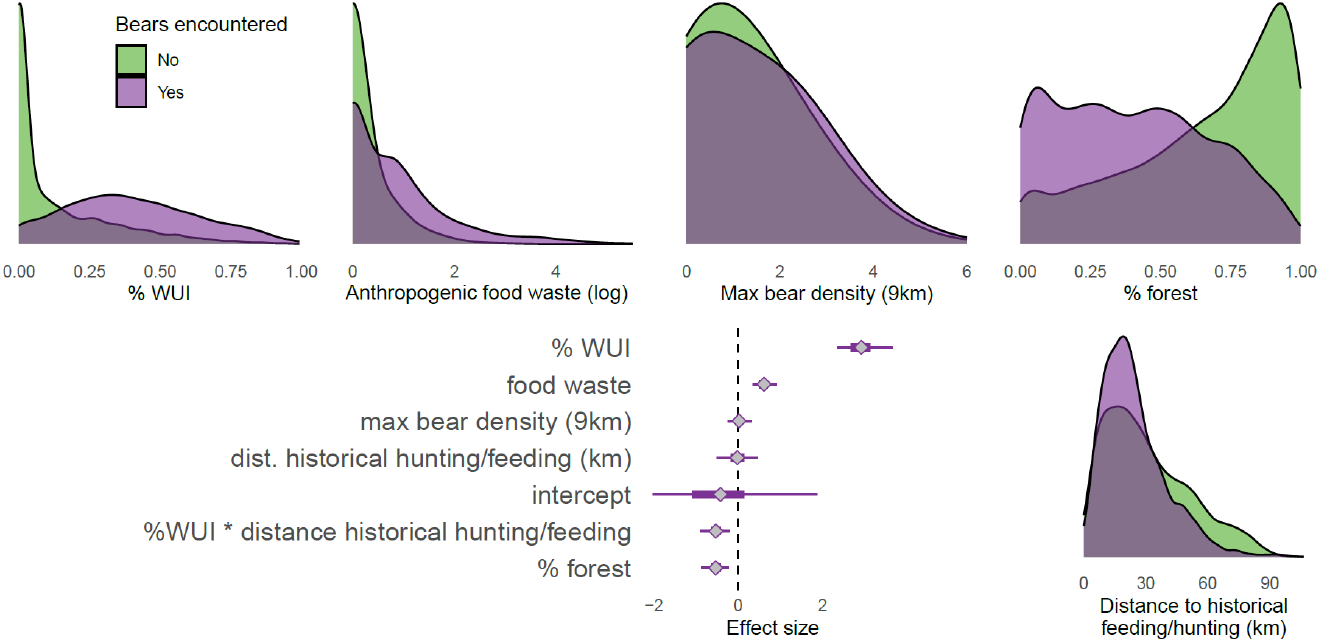
Density distribution of variables included in the occupancy model for the grid cells where bears were reported (N=1548) in purple, and for areas without bear reports in green (N=25649). Bear encounters are more frequent in areas with higher percentage of Wildland-urban-interface, intermediate forest cover, higher number of amenities, in areas with higher bear densities in the surrounding region and closer to the historical bear breeding spots. Last panel shows the relationship between the frequency of reported bears per grid cell in relation to the distance from the historical breeding facility. Central figure: Effect size (posterior mean, 50 and 95 CI) of 4 potential direct (percent WUI, percent forest cover, no. of amenities and bear density in surrounding area) and one legacy driver (distance to historical elite hunting districts) for bear occurrence in human-dominated areas. The percentage of WUI and number of sources of anthropogenic food waste indicate a strong positive relationship with bear occurrence. No effect is observed for the density of bears in the surrounding landscape or for the distance to historical hunting/feeding grounds. The percent forest cover has a negative effect on bear encounters, while the distance to the historical elite-hunting and feeding grounds has a modulating effect on the WUI.

**Figure 6A.**
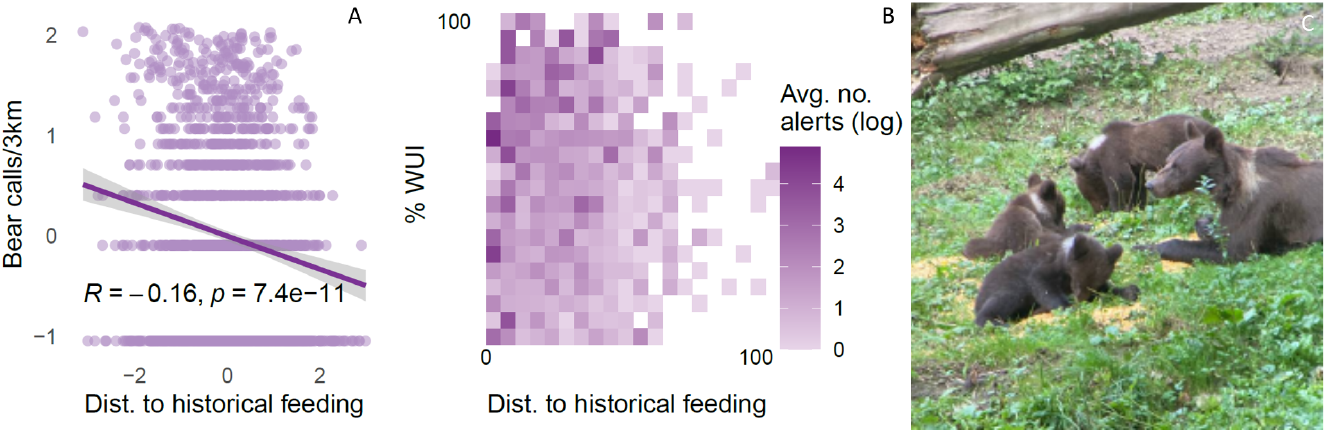
The number of bear calls per 3x3 km grid cell is decreasing with increasing distance from historical supplemental feeding locations (Spearman’s Rho -0.16, p < 0.001, variables are transformed using a Yeo-Johnson normalization). B: Interaction between % WUI (y-axis) and Distance to historical elite hunting districts (x-axis) Darker colors indicate higher average number of calls. C: Female and 3 cubs feeding on supplemental corn feed at a location is situated within < 3 km of historical supplemental feeding locations.

### Legacy effects for human-wildlife interactions

In Romania, supplemental feeding was historically tied to bait hunting and elite hunting drives (Mooney 1991; Crisan 2010). After the fall of communism, hunting became more broadly accessible (Salvatori et al. 2002), which slowed population growth (Fig 2) and likely maintained a landscape of fear (Lodberg-Holm et al. 2019; Gaynor et al. 2019). While legally approved and practiced in some parts of the world (Candler et al. 2019; Bischof et al. 2008; Lafferty et al. 2024), bait hunting remains ethically controversial (Carter and Linnell 2016) and reinforces habituation in target and non-target species (Selva et al. 2017; Penteriani et al. 2018). While globally bait hunting can generate revenue for management and conservation (Salvatori et al. 2002; Stăncioiu, Micu, et al. 2019),it also incentivizes continued feeding, which in turn can lead bears to selectively exploit locations where food is predictably provided (Powell et al. 2022; Kavčič et al. 2015), irrespective of the species targeted by feeding (Selva et al. 2017). Furthermore, supplemental feeding—whether diversionary, for bait hunting, or for wildlife watching—causes bears exposed to predictable food sources to reduce their home ranges and alter feeding habits or space use (Penteriani et al. 2021; Kavčič et al. 2015). Historically, these behavioral effects were offset by hunting pressure maintaining a landscape of fear (Lodberg-Holm et al. 2019; Gaynor et al. 2019). However, Romania’s hunting bans in 2007 and 2016 reduced perceived risk from people, allowing bears to expand into anthropogenic spaces, similar to patterns observed in Alaskan brown bears (Wheat and Wilmers 2016).As the landscape of fear dissolved and the number of habituated bears increased, conflicts, damages, and economic losses increased (Pop, Dyck, et al. 2023; Pop, Gradinaru, et al. 2023; Bautista et al. 2017; Cimpoca and Voiculescu 2022), alongside public distrust in policies (Neagu et al. 2022; Neagu and Rozylowicz 2024) and negative perceptions of the species (Stăncioiu, Dutcă, et al. 2019). Over time, these feedback loops can reinforce conflict, increase anthropogenic resistance (Ghoddousi et al. 2021) and contribute to rising biophobia (Soga and Gaston 2016). Our results provide further evidence that historical management legacies persist long after political change, continuing to shape wildlife behavior and the prospects for coexistence (Carter and Linnell 2016).

Global studies on bear behaivour suggest that increased bear densities can alter habitat use, particularly among females with cubs (Uzal et al. 2022), making high density a potential mechanism for food conditioning (Elfström et al. 2014). Our data did not support the hypothesis that high bear density in core habitats drives bear occurrence in urban or peri-urban areas, even though conspecific pressure is often cited as a factor in bears exploiting human food sources (Gunther and Wymann 2008; Powell et al. 2022). Romania has one of the highest bear densities globally, and this may mask effect sizes overall. Furthermore, to best assess this effect, we would need to include both information on change in density over time, as well as sex-ratios, both of which were not available to us for this study. We examined the proportion of RO-ALERT messages reporting females with cubs, cubs alone, or young bears and found that less than 10% of 10,672 records mentioned cubs (5.7%) and or females (3.6%), but we caution these values are unlikely to indicate true demographics, because reports heavily depend on observer species knowledge and local conditions, introducing large reporting biases.

### Sustainability implications

With the largest brown bear population in Europe west of Russia, Romania holds the potential to become a poster child for ecologically and socially sustainable bear conservation—but this requires confronting historical management legacies. Ongoing debates about conservation and population management could provide opportunities to break reinforcing cycles of habituation induced by supplemental feeding and waste management, and focus on emphasizing behavioral adaptation in both bears and humans to foster coexistence. In 2016 targeted feeding for hunting was banned in Romania, and since November 2025 also supplemental feeding for tourism has been banned. However, we suggest that further management actions could include phasing out non-target supplemental feeding (esp corn) and securing food-waste sources to reduce bears’ attraction to people. Lowering encounter rates also depends on re-establishing a landscape of fear through selective removals, hunting, or non-lethal deterrents (Sarmento 2024). Equally, addressing human habituation to bears—such as feeding or close approaches—requires targeted education in schools and popular media.

## Methods

### Study region

We studied the entire range of the brown bear (Ursus arctos) in Romania, focusing on the Romanian Carpathians—the Romanian portion of the Carpathian Ecoregion (135,000 km^2^, 57% forested), dominated by deciduous and mixed forests (Fagus sylvatica, Quercus spp.) (Munteanu et al. 2022). Romania has a population of approximately 19.5 million, with 54% living in urban areas and the remainder in smaller rural settlements (Pop, Gradinaru, et al. 2023). The country harbors the largest brown bear population in Europe outside Russia, estimated at 10,419–12,770 individuals (Ministerul Mediului, Apelor si Padurilor 2025) and one of the highest brown bear densities worldwide.

Following direct persecution and two world wars, Romania’s bear population dropped to ∼860 individuals after WWII (Quammen 2003). Hunting was banned for the general public between 1963 and 1989 but remained permitted for political elites, reflecting Nicolae Ceauşescu’s passion for hunting and use of wildlife to display political power (Quammen 2003)(Salvatori et al. 2002). During this time Romania was divided in some 2156 hunting districts, with 122 reserved for elite hunting only, and where supplemental feeding was practiced to reduce winter mortality, increase populations, individual fitness and provide predictable locations for bait-hunting (Crisan 2010; Georgescu 2003; Mooney 1991). We note that although not documented here, supplemental feeding likely occurred in other areas too. Around these historical elite-hunting management areas, ∼500 districts were designated for strict species protection. Bear numbers increased from ∼4,000 in 1975 to ∼8,000 in 1989 (Mooney 1991), although Ceauşescu and his elites reportedly hunted a total of some 450 bears until 1989 (Dominic 2000; Georgescu 2003; Crisan 2010). After 1990, hunting was opened according to management plans, and by 2000 the bear population was estimated at 5,000–6,000 (Stăncioiu, Micu, et al. 2019). A hunting ban for population management was imposed in 2007, and only culling of human-conflict causing bears was allowed until 2024, with the exception of the year 2016 when a complete ban was active (Stăncioiu, Micu, et al. 2019). Human-bear conflicts have increased steadily since 1990, with 26 fatalities and 274 injuries reported since 1995. Comparatively, Romania is characterized by both the largest number of bear attacks on humans worldwide, and the highest brown bear density (Bombieri et al. 2019),and likely many attacks remain unreported. In 2024 a level of intervention of culling 450 was approved to prevent such conflicts. To analyze bear-human interactions and their drivers, we covered the entire study area with a 3 × 3 km grid (N = 27,197 cells) with all data reported the grid-cell level.

### Bear encounters data

We used data from the RO-ALERT system, which broadcasts emergency warnings via mobile networks under the Romanian Ministry of Internal Affairs and the Inspectorate for Emergency Situations (IGSU) (Ordonanta de urgenta privind operarea sistemului de avertizare a populatiei in situatii de urgenta “RO-ALERT” 2017). Data was pre-filtered by data providers on messages that involved brown-bears (hereafter bear-alerts). The bear-alert system was launched in late 2018, and it has since curated and disseminated wildlife encounter reports based on emergency calls, primarily from populated areas.Data providers pre-filtered messages involving brown bears (hereafter *bear alerts*). Launched in late 2018, the system collects wildlife encounter reports from emergency calls, mainly in populated areas. We retrieved 10,716 bear alerts recorded between January 2019 and December 2024. Each message included location, date, and time, and occasionally the number of individuals (2.84%). We geolocated the messages using the pandas package and OpenStreetMaps OSM in python, assigning exact coordinates when specific addresses (street, house number) were available and using commune or village centroids when addresses were missing. All locations were cross-checked against message text and broadcast area, and 623 mislocated points were manually corrected. Messages lacking sufficient detail for geolocation were excluded, resulting in 10,476 georeferenced bear encounters across 1,548 unique grid cells. For each bear-alert we extracted time of day and day of year, classifying them into seasons and crepuscular periods using the ‘suncalc’ package in RStudio (Thieurmel and Elmarhraoui 2022).

### Environmental variables

We selected environmental variables expected to influence both bear detectability and their use of urban and peri-urban areas (Table 1). To account for variation in detectability, we included time of day and day of year, as bears exhibit crepuscular activity patterns and increased movement during periods of hyperphagia in spring and fall (González-Bernardo et al. 2020). We also considered population density, touristic facility density, and road density as potential factors affecting detection. We calculated population density per grid cell based on GEOSTAT 2011 grid data (Eurostat 2011). We extracted road length and the number of touristic facilities (hotels, restaurants, pensions) per grid cell from OpenStreetMap using the OSMDownloader plugin in QGIS. We focused on five environmental variables that captured the availability of anthropogenic food sources, potential for habituation, and habitat characteristics and which we expected could affect bear use of urban spaces (Table 1). Forest cover was extracted from (Hansen et al. 2013) and the proportion of the wildland–urban interface (WUI) per grid cell was calculated to identify areas with high human–wildlife conflict potential (Schug et al. 2023; Kaim et al. 2025). We quantified the density of food-related attractants per grid cell by extracting the number of potential food source locations—restaurants, hotels, supermarkets, waste containers, and food stands—from OpenStreetMap using the OSMDownloader plugin in QGIS . To represent intra-species competition and predator avoidance, we used bear density estimates for most Romanian hunting districts (INCDS: Institutul National de Cercetare-Dezvoltare in Silvlicultura 2021). We spatially allocated the average density values for the hunting district to only forested areas, extracted them to each grid cell, imputed missing values using the ‘MuMIn’ package in R, and calculated the maximum bear density within a 9 × 9 km moving window. The size of the moving window was meant to capture the approximate size of a wildlife management unit(ca. 100km2). To assess historical legacy effects, we used a 1986 hunting district map identifying 122 elite-hunting areas (average 85km2) along with their respective buffer/protection zones, noting that all these units are located within the brown-bear range but some were managed for other wildlife. We geolocated each district’s centroid and computed the distance from each grid cell to the nearest hunting district as a proxy for potential legacy effects

**Table 1:** Variables considered in the final model, the ecological processes that it proxies the expected directionality of the relationship with bear urban and peri-urban habitat use.

| **Variable** | **Expected relationship** | **Mechanism** | **Reference** |
| --- | --- | --- | --- |
| **Detectability variables** |  |  |  |
| Time of day | + | Bears are crepuscular, most active dawn and dusk | (Ordiz et al. 2017) |
| Time of year | - | Immediately after emergence from hibernation or in the fall when fattening up | (Ordiz et al. 2017; Cimpoca and Voiculescu 2022) |
| Road density | + | Measure of effort: higher likelihood of encounters due to higher human presence | (E. D. Brown and Williams 2019) |
| Population | + | Measure of effort: higher likelihood of observers being present and making calls | (E. D. Brown and Williams 2019) |
| Tourism | + | Measure of effort: higher likelihood of observer presence | (E. D. Brown and Williams 2019) |
| **Occupancy variables** |  |  |  |
| Maximum bear density in surrounding 9km area | + | Competition and predation refuge | (Elfström et al. 2014; Pop, Dyck, et al. 2023) |
| Anthropogenic food waste sources | + | Food conditioning and habituation | (Elfström et al. 2014; Powell et al. 2022; Smith et al. 2005) |
| Forest cover | - | Habitat & natural food availability | (González-Bernardo et al. 2020; Elfström et al. 2014; Pop, Dyck, et al. 2023) |
| Wildland-urban-interface (WUI) - proxy for orchards, yards and subsistence farming | + | Food availability from small farms, orchards, bee hives | (Kaim et al. 2025; Schug et al. 2023; Blecha et al. 2018) |
| Distance to historical elite hunting and supplemental feeding areas | - | Management legacy, potentially related to habituation through continued supplemental feeding | (Penteriani et al. 2021; Selva et al. 2017; Elfström et al. 2014) |

### Occupancy models

To investigate the drivers of brown bear presence in urban and peri-urban areas, we used a Bayesian hierarchical framework and a single-species spatial occupancy model implemented in the e spOccupancy package in R (Doser et al. 2022). Our data reflects structural characteristics typically used to assess human–wildlife interactions—such as citizen science observations, damage reports, emergency calls, or compensation claims (Ravenelle and Nyhus 2017; Pop, Gradinaru, et al. 2023). Occupancy models offer a robust analytical approach for such data because they incorporate detection processes (van Strien, van Swaay, and Termaat 2013; Altwegg and Nichols 2019) and control for spatial autocorrelation in addition to assessing the drivers of a species habitat use (Doser et al. 2022; Pflüger et al. 2024). We examined all variables for collinearity using Spearman’s rho and retained only predictors that did not exceed a correlation of 0.7 (Dormann et al. 2007). We log-transformed the waste and tourism variables, and normalized all predictors to mean 0 and standard deviation 1 using the ‘bestNormalize’ package in RStudio (Peterson 2023). Assuming a closed bear population between 2019 and 2024, we compiled annual presence–absence data across 1,533 grid cells with confirmed bear presence and 3,467 randomly selected absence cells located within the species’ known range (Chapron et al. 2014), resulting in a total sample of 5,000 grid cells. For each year and cell we summarized the number of bear detections, the median minute of the day and day of the year in the most frequent time and season, and the total number of bear alert messages. For absence cells, we randomly assigned time of day and day of year values. In a hierarchical modelling approach, detection probability was modelled as a function of time of day, day of year, human population, tourism potential, and road density (Table 1). Further, bear occurrence probability incorporated the detection processes and additionally included the percentage of wildland–urban interface (WUI), forest cover, potential food waste sources, local maximum bear density (9 × 9 km), and distance to historical supplemental feeding sites (Table 1). Among possible interactions, only the interaction between WUI and distance to feeding sites showed a significant effect and was retained in the final model. The final mode was calibrated using five Markov Chain Monte Carlo (MCMC) chains with 2,000 iterations, a burn-in of 2,000, and a thinning rate of 20.

We used the Bayesian posterior distributions of the coefficient estimates to assess the effects of predictors on bear site occupancy. We considered variables to have a strong effect when the 95% credible interval (CI) of the estimate did not overlap zero, and a weak effect when the 50% credible interval (CI) did not overlap zero. To evaluate model convergence, we calculated Gelman–Rubin diagnostic statistics (Rhat < 1.1) and effective sample sizes (ESS > 100) for all variables (Doser et al. 2022). We obtained 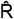 values below 1.1 and ESS values above 300 for all parameters, and visually inspected trace-plots, all indicating good model convergence and mixing. To assess model fit, we conducted posterior predictive checks using the Freeman–Tukey statistic and calculated the Bayesian p-value (Doser et al. 2022; Pflüger et al. 2024), which was 0.65, suggesting very good model fit. To further explore legacy effects of historical supplemental feeding sites, we analyzed the relationship between the number of bear-alert messages per grid cell and the distance to historical elite hunting districts, transforming both variables using a Yeo-Johnson transformation and applying a simple linear model applied only to cells with reported bear presence. We conducted all data analyses in R (R Core Team, 2020) using the packages tidyr, dplyr, sf, suncalc, terra, bestNormalize, and MuMIn for data preparation, spOccupancy for modelling and diagnostics, and ggplot2 and cowplot for visualization.

## Author contributions

CM conceptualized the study analyzed and interpreted the data and wrote the manuscript, NMD and SR compiled and analyzed data, all authors interpreted the results, IS and CM acquired funding, all authors revised and agreed with the manuscript.

## Acknowledgements

We thank Stephanie Bethmann, Bogdan Boghian, Alex Girdan, Johannes Kamp, and Vasile Tatar for valuable comments at various stages of this study. We are grateful to IGSU Romania esp. Nicusor Ionita for data support. We gratefully acknowledge support from the German Science Foundation (DFG), Research Training Group ConFoBi (GRK 2123/1 TPX).

